# NoButter: An R package for reducing transcript dispersion in CosMx Spatial Molecular Imaging Data

**DOI:** 10.1101/2024.11.25.625243

**Authors:** Béibhinn O’Hora, Roman Laddach, Rosamond Nuamah, Elena Alberts, Isobelle Wall, Joseph Bell, David A Johnston, Sonya James, Jeanette Norman, Mark G. Jones, Ciro Chiappini, Anita Grigoriadis, Jelmar Quist

## Abstract

**Motivation:** Advances in spatial transcriptomics technologies at single-cell resolution have high-lighted the need for innovative quality assessment approaches and improved analytical tools. Imaging-based spatial transcriptomics technologies, such as the CosMx Spatial Molecular Imager (SMI), provide the location and abundance of transcripts through multifocal imaging. Optical sections (or Z-slices) form a Z-stack that represents the tissue depth. Transcript dispersion can be observed across these Z-slice and introduce considerable levels of technical noise to the data that can negatively impact downstream analysis.

**Package Functionality:** NoButter is an R package designed to evaluate transcript dispersion in CosMx SMI spatial transcriptomics data. Using the raw data, the transcript distribution is assessed for each Z-slice of a Z-stack across multiple fields of views (FOVs). To systematically identify transcript dispersion, the percentage of transcripts located outside cell boundaries is calculated. Z-slices exhibiting high levels of transcript dispersion can be excluded, while high-confidence transcripts are preserved.

**Usage Scenario:** To demonstrate the functionalities of NoButter, spatial transcriptomics data was generated using the CosMx SMI for lymph node tissue, a lung sample, and two triple-negative breast cancers (TNBCs). Use cases illustrate substantial transcript dispersion in optical planes closer to the glass slide. In these Z-slices, on average, an additional 10% of the transcripts were discarded using NoButter. Cleaning such Z-slices with high dispersion rates reduces technical noise and improves the overall quality of the spatial transcriptomics data.

**Availability:** The package can be accessed at https://github.com/cancerbioinformatics/NoButter.

## Introduction

The field of spatial transcriptomics is advancing rapidly, as evidenced by the growing number of sequencing- and imaging-based technologies (Rao, et al., 2021; Tian, et al., 2023). Spatial transcriptomics is used to quantify the expression of transcripts within individual cells of a tissue thereby enhancing our understanding of the molecular phenotypes that shape its architecture. Due to the novelty of these technologies, comprehensive approaches to assess the data quality are lacking.

In imaging-based technologies, such as the CosMx Spatial Molecular Imager (SMI) (NanoString, Seattle, United States) or Xenium (10x Genomics, Pleasanton, United States), the precise localisation of transcripts is critical to accurately quantify the expression at single-cell resolution and is ultimately crucial for ensuring the reliability of any downstream analyses, including the assignment of cell types. Currently, quality control procedures for imaging-based spatial transcriptomics technologies largely follow those used in single-cell RNA-sequencing. However, imaging-based spatial transcriptomics would greatly benefit from tailored procedures to address their unique challenges.

Transcripts detection in imaging-based spatial transcriptomics is done by multifocal imaging. The resulting 3D coordinate system consists of the x- and y-axis, representing tissue width and length, and the z-axis, which directly relates to tissue depth. The CosMx SMI collects optical Z-slices at an 0.8μm interval (He, et al., 2022). Lower Z-slices (e.g. Z_0_, Z_1_) correspond to the apical plane, which is closer to the imaging objective. In contrast, higher Z-slices (e.g. Z_7_, Z_8_) capture basal planes, positioned closer to the glass slide on which the tissue rests (Figure 1A). Transcript dispersion can be defined as the movement of transcripts away from their original location. Assuming an even distribution of cells within the tissue, one would expect a similar number of transcripts in each Z-slice of a given Z-stack. However, upon inspecting data from various tissue types and CosMx SMI instruments, we found this is not always the case. In data from a lymph node, transcripts were more abundant in Z_1_-Z_2_ compared to Z_3_-Z_7_ (Supplementary Figure S1A). In contrast, in a lung and a triple-negative breast cancer (TNBC) (TNBC01) sample, transcripts were more abundant in Z_6_-Z_7_ compared to Z_-2_-Z_5_ (Supplementary Figure S1B and S1C), whereas in another TNBC (TNBC02), no difference was observed (Supplementary Figure S1D). An increase in the percentage of transcripts located outside cell boundaries was also noted in Z-slices closer to the glass slide (i.e. Z-slices with a higher index) (Figure 1B and Supplementary Figure S2). Both these findings suggest transcript dispersion has occurred, which contributes to noise in the data that can negatively impact downstream analysis.

**Figure 1.**
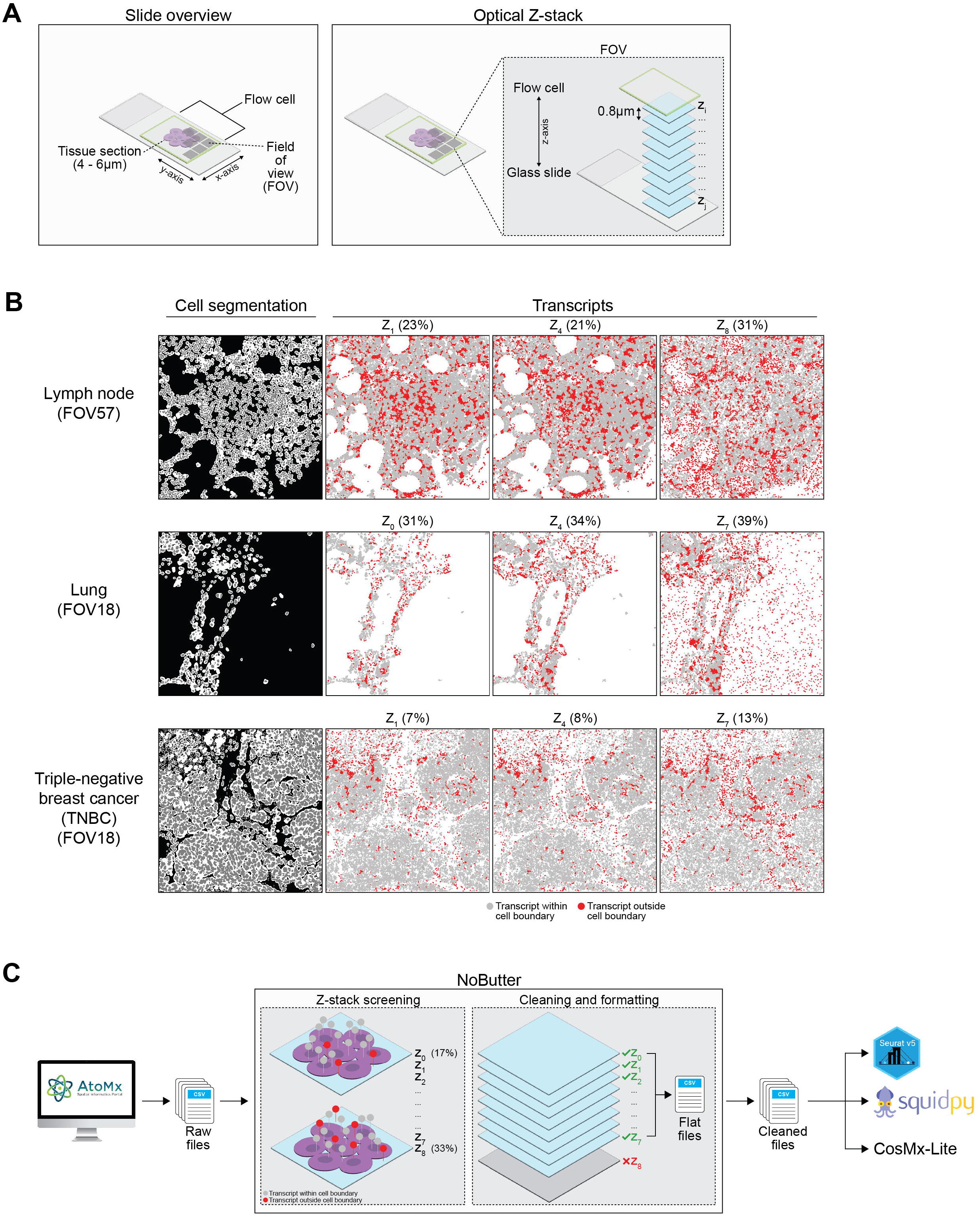
NoButter to detect and remove transcript dispersion in CosMx SMI data. **(A)** Schematic highlighting the slide layout, including tissue placement and thickness, flow cell placement, field of view (FOV) selection, and x-, y- and z-coordinates. The latter denotes the optical Z-slices that together form a Z-stack. **(B)** Examples of transcript dispersion in lymph node tissue (top, from King’s College London), a lung sample (middle, University of Southampton) and a triple-negative breast cancer (TNBC) (TNBC02) (bottom, NanoString’s Amsterdam CX Lab). Transcripts in red are located outside cell boundaries. For each tissue, three representative Z-slices are shown. The percentage of transcripts located outside the cell boundary is noted. **(C)** Schematic representation of NoButter.

To address this challenge, we developed NoButter, an R package specifically designed to assess transcript dispersion in CosMx SMI data. The package offers tools to visualise transcript distribution across Z-slices and FOVs to facilitate the detection of transcript dispersion, includes functions to discard transcripts in specific Z-slices, and to create the necessary files for downstream analysis (Figure 1C).

### Package Functionality

NoButter is created to effectively clean and preprocess CosMx SMI data. After exporting raw data from the AtoMx Spatial Informatics Platform (SIP), the package can import the contents of the ‘*AnalysisResults’* folder. This folder contains raw transcript information, such as the x- and y-coordinates, the Z-slice and the transcript type (i.e. endogenous, system or negative control), which are crucial to assess the quality of the data.

The package offers various functions to inspect raw transcript distribution. To systematically detect the presence of transcript dispersion, the percentage of raw transcripts occurring outside the cell boundaries within a given FOV should be assessed. Because transcript dispersion detection relies on the localisation of the transcripts in relation to the cell boundaries, it can be challenging to detect, particularly in FOVs encompassing densely packed cells. Therefore, it is recommended to assess FOVs that are located at the edge of the tissue section, cover gaps or breaks present in the tissue section or are positioned in regions with a distinct tissue architecture (Supplementary Figure S3).

After filtering the raw transcripts, a new transcript file can be reconstructed. The package enables the cleaning of the accompanying metadata and polygon files to reflect any necessary changes made in the transcript file. Finally, the package can consolidate these data into a set of ‘flat files’ that are compatible with various analytical tools, such as Seurat (Hao, et al., 2024), Squidpy (Palla, et al., 2022), Giotto (Dries, et al., 2021) and CosMx-Lite (O’Hora, et al., 2024).

### Usage Scenario

The applicability and functionality of NoButter are demonstrated on CosMx SMI data obtained from a lymph node, lung and two TNBCs. Slide preparation was conducted following NanoString guidelines (NanoString, 2023). The lymph node data was acquired at King’s College London, the lung at the University of Southampton and both TNBCs at NanoString’s Amsterdam CX Lab. Cell segmentation was performed in the AtoMx SIP, utilising an adaption of Cellpose (Pachitariu and Stringer, 2022). Raw transcript files were exported from AtoMx SMI using a custom export function (Griswold and Zhao, 2023).

For the lymph node, 19,132,400 transcripts were detected across 63 FOVs and 10 Z-slices. Only 4.23% of all transcripts were in Z_0_ and 3.50% in Z_8_. In Z_9_, only 261 transcripts were detected. The remainder of the transcripts (92.15%) were detected in Z_1_ to Z_7_. In the lung sample, 18,451,364 transcripts were detected in 206 FOVs and 13 Z-slices. 90.07% of the transcripts were in Z_-1_ to Z_6_. In Z_-4_ to Z_-2_, 1.71% of all transcripts were detected; in Z_7_ and Z_8_, this was 8.12%. In TNBC01, 3,003,665 transcripts were detected across 13 FOVs and 14 Z-slices. For TNBC02, 4,527,301 transcripts across 18 FOVs and 10 Z-slices were detected. In both samples, the majority (97.00% and 98.50% in TNBC01 and TNBC02 respectively) of the transcripts were detected in Z_0_ to Z_7_.

In the Z-slices closest to the glass slide (i.e. those with a high index), tissue structure became less defined (Figure 1B and Supplementary Figure S2), and a higher percentage of transcripts were detected outside of cell boundaries. This phenomenon was particularly pronounced in FOVs located at the tissue edge, or in regions with gaps or breaks in the tissue. In contrast, this was less pronounced in densely packed tissue regions (Supplementary Figure S3). These observations suggest that certain in some Z-slices, transcript dispersion is more abundant, necessitating data cleaning.

After assessing the localisation of raw transcripts, those detected outside cell boundaries and transcripts flagged as system controls were discarded, leaving 14,786,502 (77.29%) transcripts in the lymph node, 10,871,694 (58.92%) in the lung, 2,532,656 (84.32%) in TNBC01 and 3,990,287 (88.14%) in TNBC02. In addition, 1,651,132 (8.63%), 810,532 (4.39%), 390,278 (12.99%) and 495,191 (10.94%) of all transcripts were removed after Z-stack screening and cleaning using NoButter. These high-confidence transcripts were used to generate a new expression matrix and to update the metadata and polygons files, for further downstream analysis.

## Conclusion

NoButter offers a comprehensive set of functions to detect, quantify and mitigate the effects of transcript dispersion in CosMx SMI data. Its applicability and functionality are demonstrated using data from different human samples, showing how uneven transcript distribution in complex datasets can be identified and corrected. By removing these transcripts, particularly those in noisy Z-slices, NoButter improves the overall data quality and reliability.

NoButter is freely available on GitHub, complete with example data and a detailed tutorial, making it highly accessible to the research community. Ongoing development will expand its compatibility with other imaging-based spatial transcriptomics technologies.

### Package and Data Availability

NoButter can be accessed at https://github.com/cancerbioinformatics/NoButter. The repository includes example data and a comprehensive tutorial to demonstrate how to use the package. The lymph node data used in this article is publicly available. The lung and triple-negative breast cancer data can be made available upon request.

## Acknowledgements

We thank James Rosekilly and Dr Cheryl Gillett from the King’s Health Partners Cancer Biobank. The research ethics approval for the human lymph node and triple negative breast cancer samples was granted by the local research ethics committee (KHP Cancer Biobank REC ref 18/EE/0025). We would also like to thank Dr Trieu My Van for her support at NanoString’s Amsterdam CX Lab (Netherlands).

## Funding

This work was supported by Cancer Research UK City of London Centre Award (CTRQQR-2021\100004) [to A.G. and J.Q.], the National Institute for Health Research (NIHR) [to B.H], the Medical Research Council (MRC) (MR/X012476/1) [to J.Q., C.C., and A.G.], Breast Cancer Now (147KL-Q3) [A.G.]. We would also like to acknowledge funding from MRC (MC_PC_MR/Y002989/1) [to J.B., D.A.J., S.J., J.N. and M.G.J.], Asthma+ Lung UK (SRG19\100001) [to J.B., D.A.J., S.J., J.N. and M.G.J.] and the NIHR Southampton Biomedical Research Centre.

## Figure Legends

**Supplementary Figure S1.**
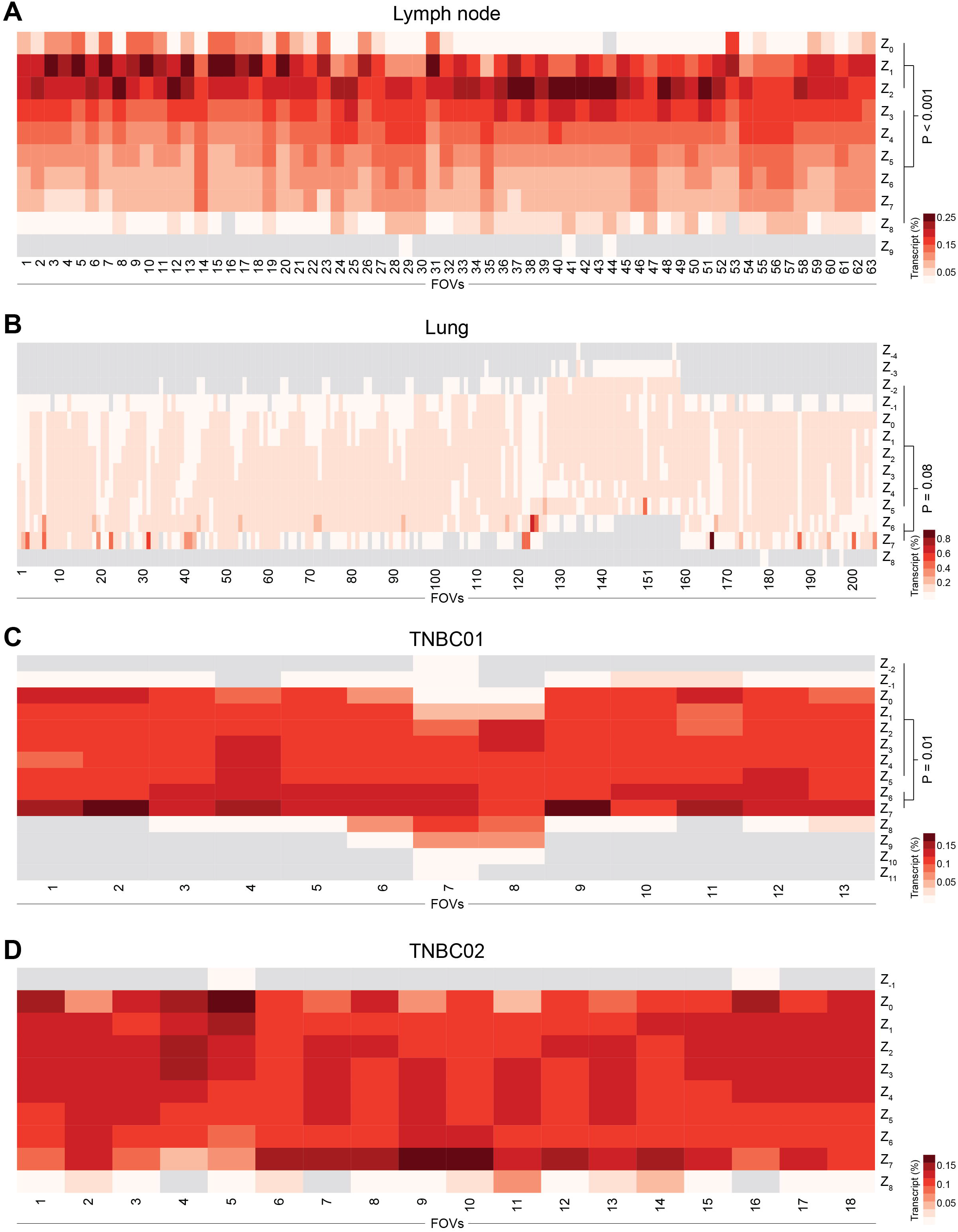
Heatmap representing the transcript distribution across Z-slices, normalised by FOV, in **(A)** lymph node tissue, **(B)** a lung sample, **(C)** TNBC01, and **(D)** TNBC02.

**Supplementary Figure S2.**
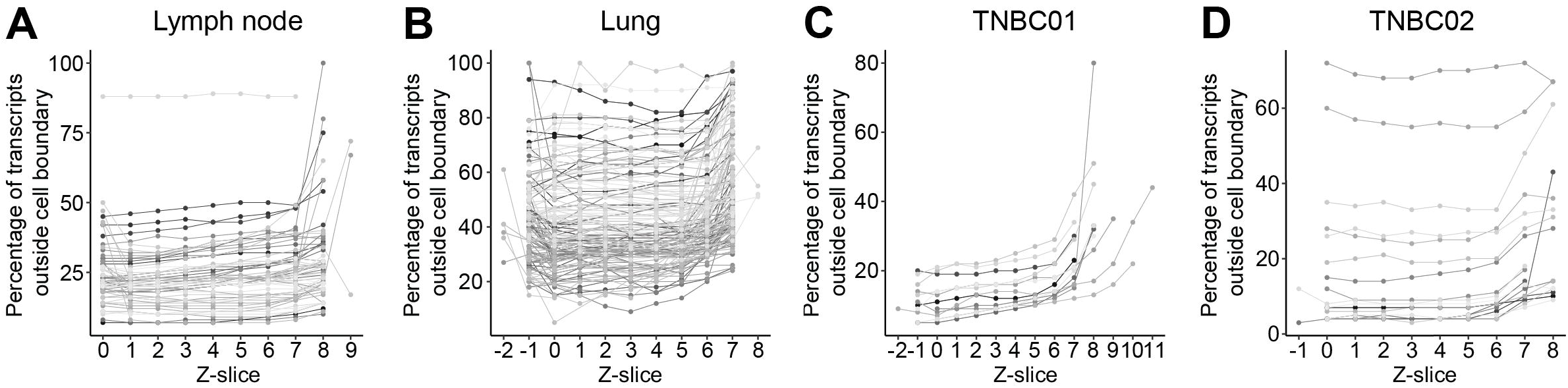
Line plot of the percentage of transcripts outside cell boundaries across Z-slices in **(A)** lymph node tissue, **(B)** a lung sample, **(C)** TNBC01, and **(D)** TNBC02. Each line represents an FOV.

**Supplementary Figure S3.**
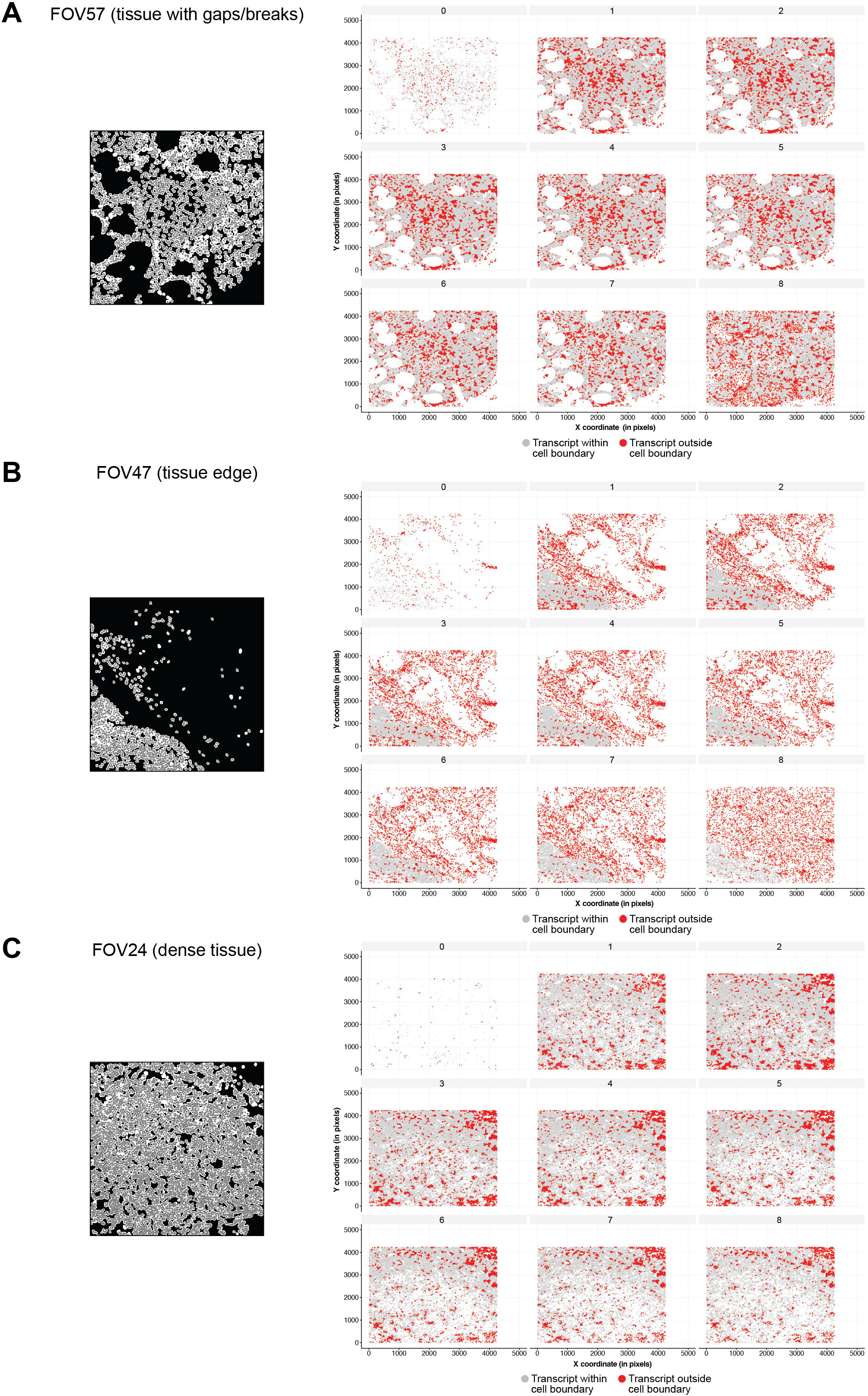
Examples depicting differences in the detection of transcript dispersion in **(A)** FOVs encompassing tissue with gaps and/or breaks, **(B)** FOVs placed at the tissue edge, and **(C)** FOVs in dense tissue.

## Notes

### Competing Interest Statement

The authors have declared no competing interest.

